# Comparative analysis of Synthetic Physical Interactions with the yeast centrosome

**DOI:** 10.1101/514166

**Authors:** Rowan S M Howell, Attila Csikász-Nagy, Peter H Thorpe

## Abstract

The yeast centrosome or Spindle Pole Body (SPB) is situated in the nuclear membrane, where it nucleates spindle microtubules and acts as a signalling hub. Previously, we used Synthetic Physical Interactions to map the regions of the cell that are sensitive to forced relocalization of proteins across the proteome [Berry et al., 2016]. Here, we expand on this work to show that the SPB, in particular, is sensitive to the relocalization of many proteins. This work inspired a new data analysis approach that indicates that relocalization screens may produce more growth defects than previously reported. A set of associations with the SPB result in elevated SPB number and since hyper-proliferation of centrosomes is a hallmark of cancer cells, these associations point the way for the use of yeast models in the study of spindle formation and chromosome segregation in cancer.

## 2 Introduction

Microtubule Organising Centres (MTOCs) are critical to the process of chromosome segregation in eukaryotes; abnormalities in the structure or number of centrosomes is strongly associated with human cancer [Nigg, 2006]. In *S. cerevisiae*, the MTOC is the Spindle Pole Body (SPB). The SPB differs from metazoan centrosomes in its structure and in the fact that it remains embedded in the nuclear membrane throughout the closed mitosis of yeast [Fu et al., 2015]. However, despite these differences, there is significant conservation between yeast SPB proteins and human centrosomal proteins [Jaspersen and Winey, 2004], making the yeast SPB a relevant model of MTOCs.

Beyond their roles in microtubule nucleation, SPBs are thought to act as signalling hubs, with recruitment to the SPB a key step in regulation of certain signalling pathways [Fu et al., 2015, Arquint et al., 2014]. Various studies have used the strong interaction between GFP and GFP-Binding Protein (GBP) [Rothbauer et al., 2006], to test the effect of forced localization to the SPB, for example Gryaznova et al. [2016], Caydasi et al. [2017]. However, no systematic study of forced relocalization to the SPB has been performed. We used the Synthetic Physical Interaction (SPI) methodology [Ólafsson and Thorpe, 2015] to test recruitment of more than 4,000 proteins to five locations around the SPB.

Proteome-wide SPI screens have been used in the past to probe the regulation of the kinetochore [Ólafsson and Thorpe, 2015, 2016] and a set of 23 SPI screens was used to generate a cell-wide map of proteins sensitive to relocalization [Berry et al., 2016]. Relative to the screens of Berry et al. our analysis shows that the SPB is particularly sensitive to forcible relocalization. As a result, we found that standard methods for analysis of genome-wide screens based on Z-transformations were unsuitable to analyse these screens. Efron [Efron, 2004] suggested an approach to multiple hypothesis testing, such as genome-wide screens, based around an empirically derived null distribution which he treated as a component of a finite mixture model to calculate significance of measured results. This empirical Bayes approach is widely used to analyse gene expression data, where it is used to classify the significance of correlations between genes, see for example [Schafer and Strimmer, 2005]. We adapted this approach to the analysis of the 23 SPI screens conducted by Berry et al. [2016] as well as the five SPB SPI screens. We used the “Mclust” package [Scrucca et al., 2016] to fit bimodal normal mixture models to our data, according to an approach outlined in [Fraley and Raftery, 2002]. This approach overcomes the limitations of Z-transformations as well as providing a parameterisation to compare screens and tools to predict the rate of validation. Berry et al. [2016] concluded that only a small fraction of proteins are sensitive to forced relocalisation. Our analysis suggests that specific regions of the cell, including the SPB, are far more sensitive to forcible relocalization.

Global analysis of the SPI data shows that the SPB is sensitive to forced interactions with a variety of classes of proteins, including proteins involved in microtubule nucleation, protein transport, lipid biosynthesis and the cell cycle. Proteins that caused growth defects when recruited to the SPB originated from the nucleus and chromosomes as well as membranes and especially the endoplasmic reticulum. Although we found significant variation in individual results between regions of the SPB, the data from the SPB SPI screens was found to be more similar to each other than to the screens with other parts of the cell. Berry et al. [2016] concluded that only ~ 2% of proteins were sensitive to forcible localization, our analysis suggests that locally this may vary with some regions, such as the SPB, far more sensitive than other parts.

A result of particular interest was that tethering nuclear pore proteins to the SPB caused growth defects. A growing body of work (reviewed in Jaspersen and Ghosh [2012], Rüthnick and Schiebel [2018]), shows that the process of SPB duplication and insertion into the nuclear membrane relies on machinery usually associated with the nuclear pore. We investigated whether these forced interactions between Spc42 and nuclear pore proteins resulted in abnormal SPB number. We found that several nuclear pore proteins, as well as the SPIN (SPB Insertion Network) and some currently-unclassified membrane proteins showed evidence of SPB overduplication. The current model for SPB duplication is that it is tied to the cell division cycle through sequential activation by Cdc14 and CDK [Rüthnick and Schiebel, 2018]. Our work suggests that forced localization of nuclear pore proteins to the SPB can decouple the process of SPB duplication from the cell cycle, a finding that may suggest refinement of the current model or that SPB duplication can occur via alternative pathways. The discovery of yeast strains that can overduplicate their SPBs may be of use as a model for cancer cells, which are known to exhibit significant variation in centrosome number [Nigg, 2006].

## 3 Materials and Methods

### 3.1 Yeast strains and methods

Yeast was cultured in standard growth media with 2% (w/v) glucose unless otherwise stated. GFP strains in this study are from a library derived from BY4741 (*his*3Δ1 *leu*2Δ0 *met*15Δ0 *ura*3Δ0) [Huh et al., 2003, Tkach et al., 2012]. For each screen we constructed plasmids expressing an SPB-GBP-RFP construct and the SPB protein alone from the *CUP1* promoter; all plasmids are derived from pWJ1512 [Reid et al., 2011] and are listed in Table 2. Plasmids were constructed by gap repair either through in vivo recombination or the NEBuilder plasmid assembly tool (New England Biolabs, USA). Linear products were created through PCR with primers from Sigma Life Science and Q5 Polymerase (New England Biolabs, USA). The sequence of azurite fluorescent protein [Mena et al., 2006] was synthesized by GeneArt (ThermoFisher Scientific, UK). The sequence of all plasmids was verified by Sanger sequencing (Genomics Equipment Park STP, Francis Crick Institute and Genewiz, UK).

### 3.2 SPI screening

The SPI screening process is described in detail in Berry et al. [2016] and in Ólafsson and Thorpe [2018]. A library of GFP strains is transformed with a plasmid expressing either a fusion of a protein of interest with GBP or a control, through a mating-based method known as Selective Ploidy Ablation (SPA) [Reid et al., 2011]. The plates are repeatedly copied and grown on successive rounds of selection media until a library of haploid GFP strains with the plasmid is produced. This library is assayed for colony size, giving a readout for the fitness of a given binary fusion between the GFP strain and protein of interest. Plates were scanned on a desktop flatbed scanner (Epson V750 Pro, Seiko Epson Corporation, Japan) at a resolution of 300 dpi. All plates were grown at 30°C. All copying of yeast colonies was performed on a Rotor robot (Singer Instruments, UK).

### 3.3 Quantitative analysis of high-throughput yeast growth

Scanned images were analysed computationally to extract measurements of the colony sizes. The online tool ScreenMill [Dittmar et al., 2010] was used to perform normalisation and calculate Log Growth Ratios (LGRs) and Z-scores by comparison of experimental and control colony sizes. Two controls were used (plasmids expressing GBP or the SPB protein alone) but, as in previous studies, we found strong agreement between the two and we used an average of the two values. In some cases, the library contained multiple copies of the same GFP strain, in these cases data was aggregated by averaging. In the proteome-wide screens plates were normalised to the plate median while in the validation screens GFP-free controls were used for normalisation. LGRs were further normalised using a spatial smoothing algorithm as described in Berry et al. [2016]. Bimodal normal mixture models were fitted to the smoothed LGR data using the “Mclust” package [Scrucca et al., 2016]. Further details can be found in the supplementary materials. R scripts for data formatting and analysis are freely available at https://github.com/RowanHowell/data-analysis.

### 3.4 Bioinformatics

The GOrilla website (cbl-gorilla.cs.technion.ac.il [Eden et al., 2009]) was used to perform all gene ontology enrichment analysis. The “cluster” program (version 3.0) [Eisen et al., 1998] was used to perform hierarchical clustering of the SPI data; Java Treeview [Saldanha 2004] was used to visualize the results. Clustering was performed using either the correlation of the LGRs, minimizing the average linkage of the clusters.

### 3.5 Fluorescence microscopy

To examine localization of the SPB-GBP-RFP construct, and GFP-tagged proteins, cells were grown shaking overnight at 23°C in -leucine media, supplemented with additional adenine. They were then imaged with a Zeiss Axioimager Z2 microscope (Carl Zeiss AG, Germany), with a 63x 1.4NA oil immersion lens and using a Zeiss Colibri LED illumination system (GFP=470 nm, RFP=590 nm, azurite=385nm). Bright field images were obtained and visualised using differential interference contrast (DIC) prisms. Images were captured using a Hamamatsu Flash 4 Lte. CMOS camera containing a FL-400 sensor with 6.5 mm pixels, binned 2×2. Images were prepared with Volocity software (Perkin Elmer Inc., USA). To screen for abnormal numbers of foci in strains containing Spc42-GBP-RFP and GFP-tagged proteins, a plate of strains was prepared using the SPA methodology described above. In this assay, the Spc42-GBP-RFP plasmid (pHT11) was accompanied by a plasmid expressing Htb2-Azurite (pHT 706) with nourseothricin (*NAT*) selection. The Htb2-Azurite construct allowed for identification of the nucleus. On the same day, cells were picked from the plate and suspended in water and them imaged as described above. Dead cells were identified by a high level of dispersed fluorescence, and were excluded, as were cells with no visible fluorescence in the RFP channel.

## 4 Results

### 4.1 Synthetic Physical Interaction screens with the SPB

The budding yeast SPB is embedded in the nuclear membrane with one face, known as the inner plaque, directed into the nucleus and the other, known as the outer plaque, facing into the cytoplasm [Jaspersen and Winey, 2004] (Figure 1A). A central plaque links the inner and outer faces of the SPB and connects to a structure known as the half-bridge, which is involved in SPB duplication. In order to understand the effect of localising proteins to different parts of the SPB, we performed genome-wide Synthetic Physical Interaction screens with multiple target proteins: Nud1, Spc42, Spc72 and Spc110 N-termini and Spc110 C-terminus GBP fusions. Nud1 and Spc72 are situated in the outer plaque of the SPB; the N-terminus of Spc110 lies on the inner plaque while its C-terminus is located, with Spc42, in the central plaque [Jaspersen and Winey, 2004].

**Figure 1:**
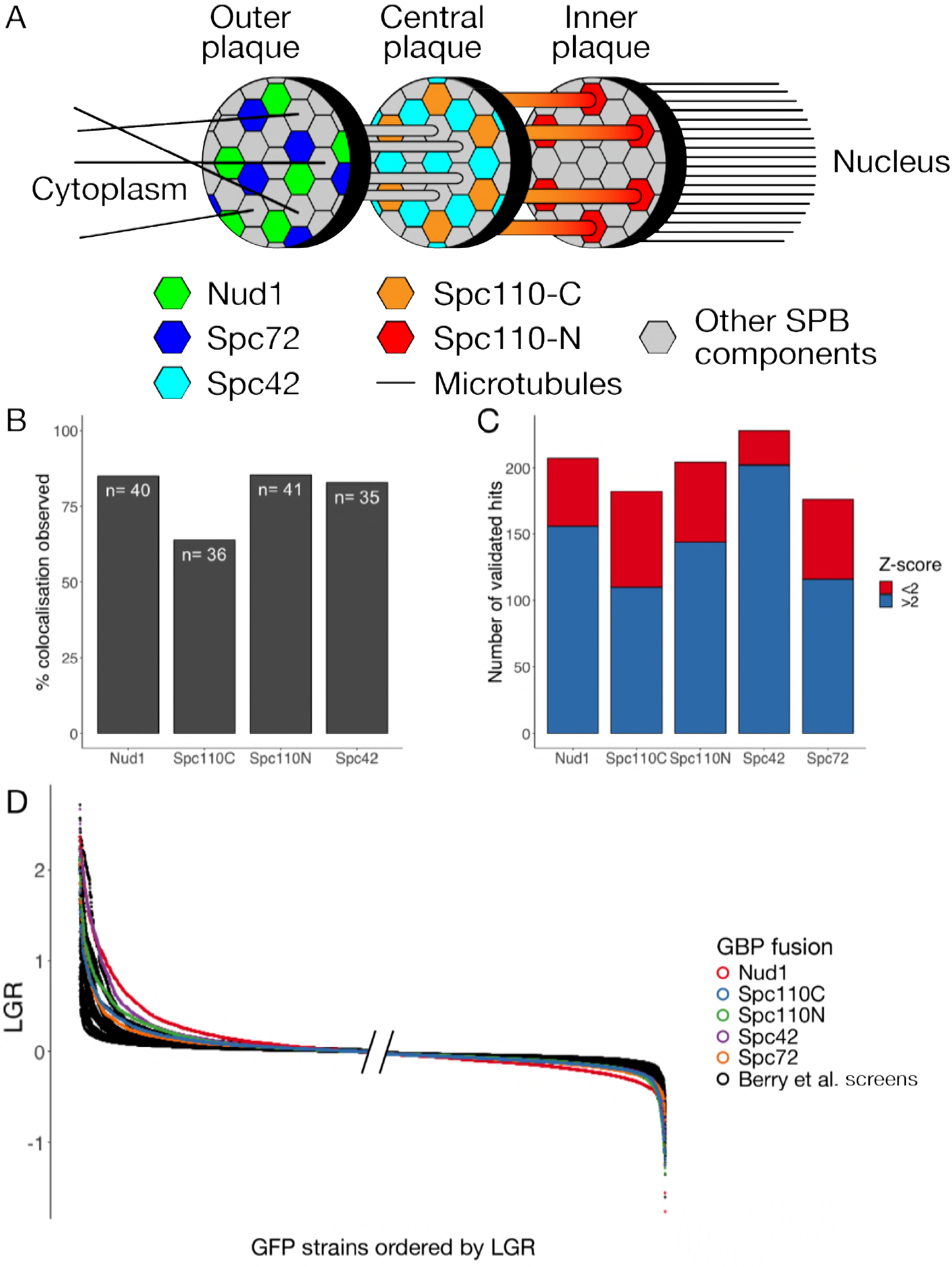
*(previous page)*: A: Structure of the SPB showing location of GBP tags used. B: Colocalization of query and target proteins in the Nud1, Spc42, Spc110C and Spc110N screens. A selection of 48 GFP strains were chosen to represent different regions of the cell and a mixture of strong and weak growth phenotypes. Each strain was judged to have either colocalization of GFP and RFP at SPB foci or not. In some cases no live cells were imaged due to slow growth, these strains were removed from analysis. The 60% − 80% colocalization observed in each screen is consistent with previous studies [Berry et al., 2016]. C: Validation of SPB SPI screens. For each GBP construct, 240 GFP strains were chosen and rescreened at higher density. These strains were considered to be validated hits if the growth defect measured was greater than a cutoff determined by GFP-free controls. In each screen, we found that strains with Z-scores less than 2 met the criteria for validation, suggesting the cutoff at a Z-score of 2 was overly restrictive. D: Ordered LGRs for each of the 5 SPB screens and 23 screens from Berry et al. [2016], this graph shows only strains present in the subset of the GFP library used in the SPB screens. The left hand side of the graph has left-justified values while the right hand side shows the right-justified values, this is because the region closest to the edges is the most informative. The SPB screens, shown in colour, are considerably seperated from the screens performed with other regions of the cell.

SPI screens aim to test the effects of forced relocalization of gene products across the genome [Ólafsson and Thorpe, 2015]. In each screen, a target gene tagged with GBP (GFP-Binding Protein) is introduced into a library of GFP strains [Tkach et al., 2012] to induce binary fusions between the target protein and the GFP-tagged query protein. Growth of colonies under these conditions is measured and an average LGR (Log Growth Ratio) between the experimental strain and two control strains is calculated, providing a measure of any growth defect caused by the artificial protein-protein interaction. Additionally, a Z-transformation is applied to assess the significance of the results. A Z-transformation assumes the data is normally distributed and uses the mean and variance of the data to transform each data point to a Z-score, which are distributed according to a normal distribution with a mean of 0 and a standard deviation of 1. This simplifies analysis, in Z-space the region (−2, 2) represents the 95% confidence interval. Similarly to genetic interactions, we say a forced association between proteins is a SPI only if this combination causes a growth defect.

We predicted that, due to the structural integrity of the SPB, the GFP-tagged query proteins were more likely to be recruited to the GBP-tagged target protein than vice versa. Fluorescence microscopy of 48 representative strains demonstrated that 60%-80% of strains viewed showed localization patterns consistent with recruitment of the query protein to the SPB (Figure 1B) in the Nud1, Spc42, Spc110C and Spc110N screens; a finding in keeping with the results of Berry et al. [2016]. Genome-wide screens often have high rates of type I errors (false positives) so we validated a selection of strains with high or, in some cases, low, negative LGRs. Validation screens were performed with 16, rather than 4, replicates of each strain and “validation” of a result was defined by a LGR exceeding a threshold set by GFP-free controls. Each of the screens identified ~ 150 strains with Z-scores greater than 2 and we validated 240 strains for each screen. The remaining strains were chosen as those just below the Z-score of 2 cutoff and “growth enhancers” - strains with Z-score less than −2, these were found mainly in the Spc110C and Spc110N screens. The growth enhancers were found not to validate frequently in either screen, these strains are likely slow growing generally, which can lead to inaccurate LGRs (see Figure 2A). We were surprised to discover that almost all of the strains with Z-scores above 2 validated and many that lay below this cutoff validated as well (Figure 1C). Furthermore, when we plotted the distribution of LGRs against LGRs from a dataset of 23 SPI screens [Berry et al., 2016], we noticed that the SPB screens generally had more, high LGR strains than other screens (Figure 1D). We hypothesized that the SPB was particularly sensitive to forced localization and that these screens identified many true hits. However this was not reflected in the number of hits according to the Z-score. As the Z-transformation is based on the assumption that data is normally distributed, it will become inappropriate when the data deviates significantly from this distribution, as we would expect in the case of a screen with many hits. Therefore, we developed a novel statistical methodology to analyze significance in SPI screens.

**Figure 2:**
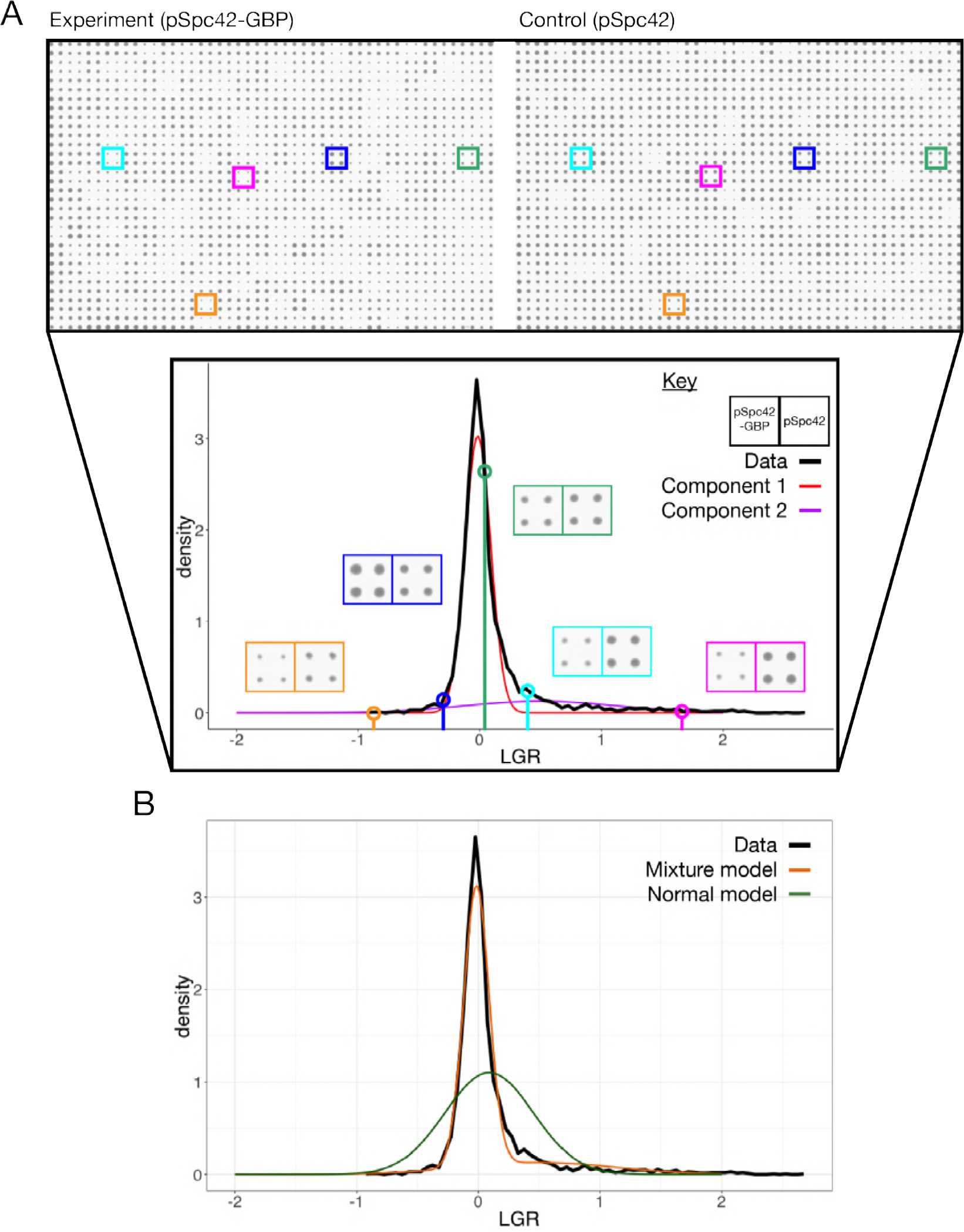
A: Schematic of the mixture model analysis of SPI screen data for the Spc42 screen. The top panel shows scans of a single library plate with the plasmid expressing either Spc42-GBP-RFP (denoted Spc42-GBP) or Spc42 (the plate with the plasmid expressing GBP alone is not shown). The lower panel shows a histogram of LGRs in the screen, with two normal components of the mixture model shown in colour. Five strains are highlighted to show the difference in colony size associated with different LGRs. Note that strains with low negative LGRs, such as that shown in orange are often the results of slow-growing GFP strains, which can register as having enhanced growth due to plate normalization and proportionally high levels of measurement error. B: Comparison of the bimodal normal mixture model and normal model of the Spc42 screen data, with the histogram of measured LGRs.

### 4.2 Mixture models are an effective model for SPI screen data

Based on the approach of Efron [2004], we developed an empirical Bayes methodology to analyze SPI screen data based on mixture models. Genome-wide screens, such as SPI screens, typically apply an experimental procedure to assign every gene in the genome a value. In yeast screens, such as SPI or yeast-two-hybrid screens, this measure often characterizes the growth of a colony. Analysis of these screens generally assumes that the distribution of colony sizes under equal conditions will follow a lognormal distribution, so that the logarithm of these sizes is normally distributed. However, when performing a genome-wide screen, we expect some small but non-zero proportion of strains to have reduced fitness and grow more slowly. We hypothesized that in certain cases, where a significant number of genes are affected, screening data will not fit a normal distribution. In a previous study, Berry et al. [2016] performed 23 SPI screens using GBP fusions in different compartments of the cell, in order to build up a map of protein localization sensitivity. We combined this dataset of screens with the SPB SPI data to assess the performance of Z-transformations in different proteome-wide screens.

We found that the LGRs are not distributed according to a normal distribution (Figure 2B). We reasoned that we could take advantage of the assumption that the data contained two distinct categories, unaffected and affected by forced localization, to develop an improved statistical model of the data. We used the “Mclust” package [Scrucca et al., 2016] to fit bimodal normal mixture models [Fraley and Raftery, 2002] to the SPI data (Figure 2). These mixture models matched the distribution of SPI data more successfully than uni-modal normal distributions (Figure 2B). We found that for 20 of the 28 screens the fitted mixture model matched our intuition of a “central” peak representing unaffected genes and a “hit” peak, shifted to the right representing genes affected by the forced interaction. The data for the remaining eight screens did not show well-defined hit peaks. An underlying assumption of our analysis is that the non-hits will be distributed according to a normal distribution, so in screens with few hits, we would expect a normal model to fit the data effectively. We interpret the failure of the mixture model to identify a well-defined hit peak in these eight screens as indicating that the screens have few hits and that therefore, in these cases, a Z-transformation would be appropriate. When present, the two overlapping, peaks in the data allows for the identification of two defined categories in the data. Component 1, or the central peak, contains genes unaffected by the interaction and is distributed normally due to noise in measurement. Component 2, or the hit peak, contains genes affected by the interaction, the shape of this distribution represents both effects of noise and the distribution of strength of real growth defects. We do not know *a priori* the shape of the distribution of interactions effects, but here we make the assumption it is gaussian.

### 4.3 Tools based on mixture models

Having determined that mixture models are a more appropriate statistical model than a normal distribution, we developed metrics to determine the significance of individual results and cutoffs to distinguish hits from non-hits. A typical approach in genome-wide screens is to calculate p-values based on a null model of the data. In the case of mixture models, identifying Component 1 as an empirical null model for the data allows for calculation of p-values, which may be adjusted for multiple hypothesis testing, for example by calculating FDR q-values [Benjamini and Hochberg, 1995]. However, in this context a more natural approach is to calculate the conditional probability of inclusion in Component 2. We define *q*(*x*) to be the probability of inclusion in Component 2 given a measured LGR of *x*. The point where *q*(*x*) = 0.5 is the point where a strain with measured LGR *x* is equally likely to be in Component 1 or 2, and is therefore a logical point to place a cutoff. We define this point as *L*_*q*,0.5_, while the point where a Z-transformation of the data has value 2 is *L*x_*Z*_. We found that *L*_*q*,0.5_ always sat below the *L*_*Z*_ but this effect was more pronounced in screens with more hits. Notably using Z-score as a cutoff limited the range of numbers of hits to 100-250. In contrast, using *L*_*q*,0.5_ as a cutoff has a dynamic range of 100-700 hits (Figure 3A). This makes the mixture model approach a more effective tool than Z-score to distinguish between screens with many or few hits.

**Figure 3:**
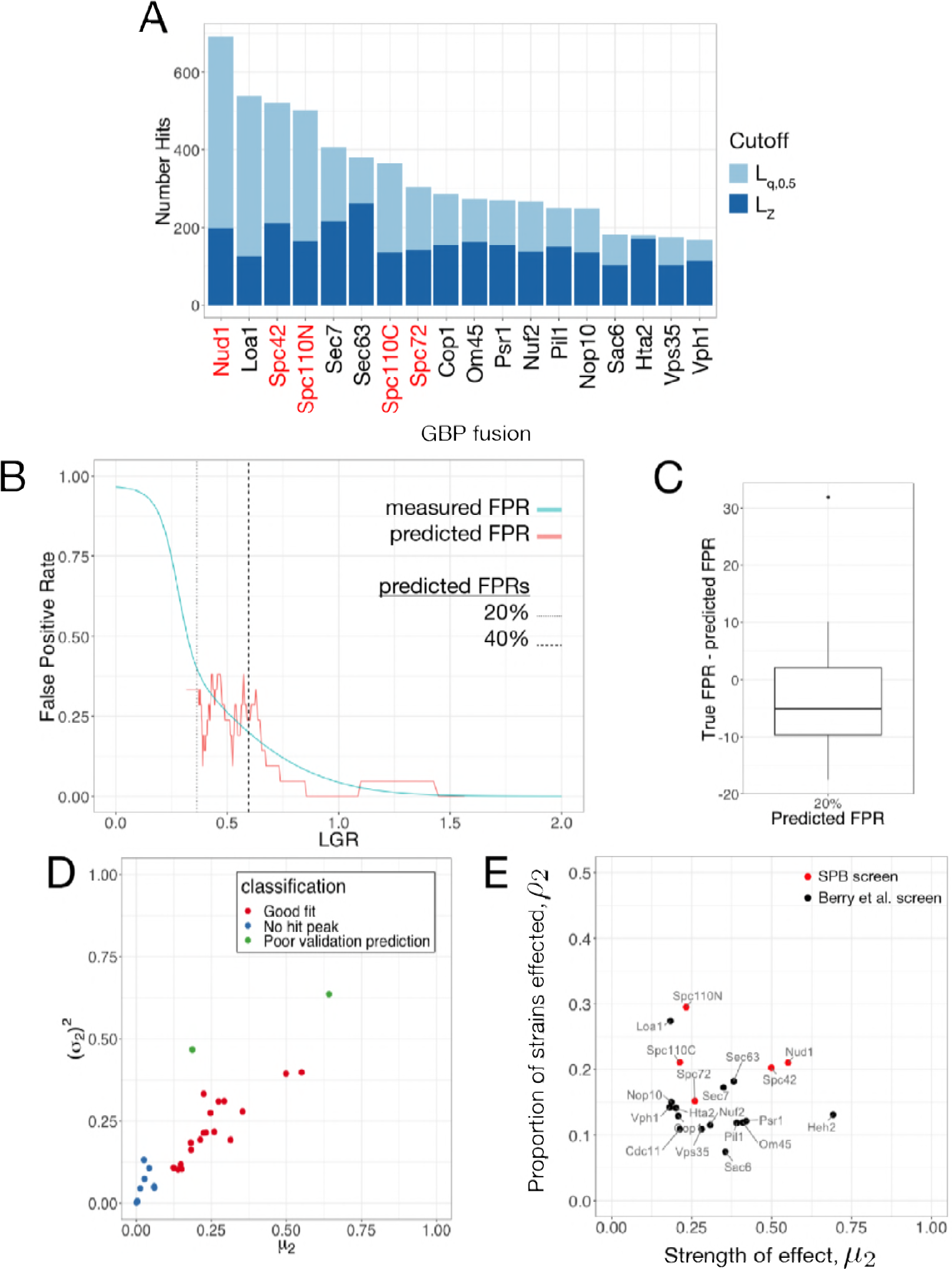
*(previous page)*: A: The number of hits by both Z-score (*L*_*Z*_) and q(x) (*L*_*q*,0.5_) cutoff for each of the screens where the mixture model was applicable. The q(x) cutoff has a higher dynamic range than the Z-score and is better able to distinguish screens with many hits. The SPB screens are among those with the greatest numbers of hits. B: FPR prediction for the Spc72 screen. The FPR for the screen was predicted from the mixture model and this prediction is overlaid with estimates of the FPR using binned data from the validation screen. In this case, the predicted FPR was reasonably accurate, although the data is quite noisy. The points where the mixture model predicts 20% and 40% FPR are indicated with a dashed line. C: Box-and-whisker plot showing the difference between measured and predicted FPR at the point where the FPR is predicted to be 20% across the screens where the mixture model was applicable. This shows some bias, with the predicted FPR generally higher than the true FPR but generally achieving an accuracy around ±10%. D: Classification of mixture model fit for each of the 28 screens analyzed. The mean *μ*_2_ and variance (*σ*_2_)^2^ of component 2 are good indicators of the success of the model with very low means or high variances indicative of the lack of a hit peak or poor validation prediction respectively. E: Classification of screen based on fitted parameters calculated using the subset of GFP strains used in the SPB screen. Each of the screens for which the mixture model fit was appropriate are plotted according to the proportion of strains affected (ρ2) and the average strength of these effects (*μ*_2_). The SPB screens Spc42 and Nud1 are positioned in the upper right portion of the graph, showing that a large proportion of proteins were sensitive to forced interaction with the SPB and these sensitivities caused significant growth defects.

The top hits from genome-wide screens are commonly validated by repeating the screen either to verify key results or to establish metrics such as the False Positive Rate (FPR). Validation is undesirable as it requires further resources and, in some cases, may not be practical, so we developed a statistical method to predict the FPR. In the validation screens, 16 replicates were used as opposed to 4 in the original genome-wide screens. Furthermore, a hit was considered to be validated if the measured LGR was considered to be significant relative to GFP-free controls. Validation is considered to be a “gold-standard” for hit verification as it corresponds closely to other assays for growth defects such as spot tests. As the *q*(*x*) cutoffs were lower than the Z-score cutoffs we were concerned that they may not be reliable indicators of validation. Indeed, most of the *q*(*x*) cutoffs lay below the 40% FPR point at which Berry et al. [2016] stopped validating. It is worth noting that results that do not validate may still be reproducible and biologically interesting despite having relatively subtle effects on growth that are difficult to distinguish from the variability in wild type growth. Therefore, we developed a method to predict the likelihood of validation. Using the fitted mixture models, we developed a metric *pV*(*x*), representing the probability of validation for a strain with measured LGR *x*. *pV*(*x*) is generally successful at predicting the rate of validation in a screen (Figure 3B). For the 20 SPI screens which were fit well by the mixture models, the validation rate of 18 of these screens was predicted well by *pV* (*x*). The other two generally had very poor validation rates in general, making any kind of validation prediction unlikely to succeed. Plotting the variance and means of Component 2 for each of the SPI screens (Figure 3D) shows that both of these screens are outliers with very high variances. Therefore, we recommend that when using this approach, great care is taken when the variance of Component 2 is high. Comparison of specific points, for example 20% FPR, shows good predictive power (Figure 3C).

### 4.4 The SPB is especially sensitive to forced relocalization

When we compared the SPI screens using SPB components with the previous screens using other structures throughout the cell, we noticed some key differences. Figure 3A shows that the SPB SPI screens are among the screens with the greatest number of hits, both by Z-score and *q*(*x*) cutoff. The fitted mixture models offer an additional way to understand this difference. Within a SPI screen, we may wish to distinguish between the case of a large proportion of strains being affected in a minor way and a smaller proportion of strains being very strongly affected. The fitted parameters *ρ*_2_ and *μ*_2_ reflect the proportion of strains affected and the severity of these effects respectively. Plotting these two parameters together therefore provides a graphical way to compare screens. This is shown in Figure 3E, where we see that the SPI screens sit in the topright region of the graph as they have high values of *ρ*_2_ and *μ*_2_. In particular, Spc42 and Nud1 produce especially strong SPIs (high *μ*_2_), while the Spc110 screens produce weaker SPIs but with many different strains (high *ρ*_2_). Spc72 is more midrange, possible reflecting the fact that in the S288C background, *SPC72* is a non-essential gene [Giaever et al., 2002]. Notably, Loa1 has a high value of *σ*_2_, Heh2 has a high value of *μ*_2_ and Sec7 and Sec63 sit near to Spc42 and Nud1. All four of these proteins localize to the ER, golgi or nuclear membrane, suggesting that these regions specifically may be the most sensitive to forcible relocalization.

We used hierarchical clustering to compare the SPB screens to the other SPI screens in the dataset (Figure 4). The data was clustered both vertically (by GFP strain) and horizontally (by screen). Clustering by screen shows the five SPB screens are more similar to each other than to other screens in the dataset, suggesting there is a characteristic set of proteins that are sensitive to forced localization to the SPB. Clustering the data by GFP strain identifies clusters of biologically related proteins with similar profiles of localization sensitivity, as previously reported [Berry et al., 2016]. Clusters of proteins with SPIs with the SPB group together, for example, the fatty acid elongases Elo1, Elo2 and Elo3; and the two paralogs of HMG-CoA reductase Hmg1 and Hmg2. These clusters also link together members of protein complexes such as the ER membrane protein complex (EMC) and oligosaccharyltransferase complex (OST). The clustering identifies a group of proteins that appear to enhance growth when forced to interact with both termini of Spc110. This group is not significantly enriched for any GO terms, however, it does include Mad2, consistent with the idea that partial Mad2 perturbation may accelerate cell cycle progression [Barnhart et al., 2011]. However, we found that growth enhancers were unlikely to reproduce their behaviour in validation screens. The vertical clustering also identifies a collection of proteins that are sensitive to forcible relocalization to all or most parts of the cell. This group of proteins, known as “frequent flyers”, are enriched for transcription factors and nuclear proteins.

**Figure 4:**
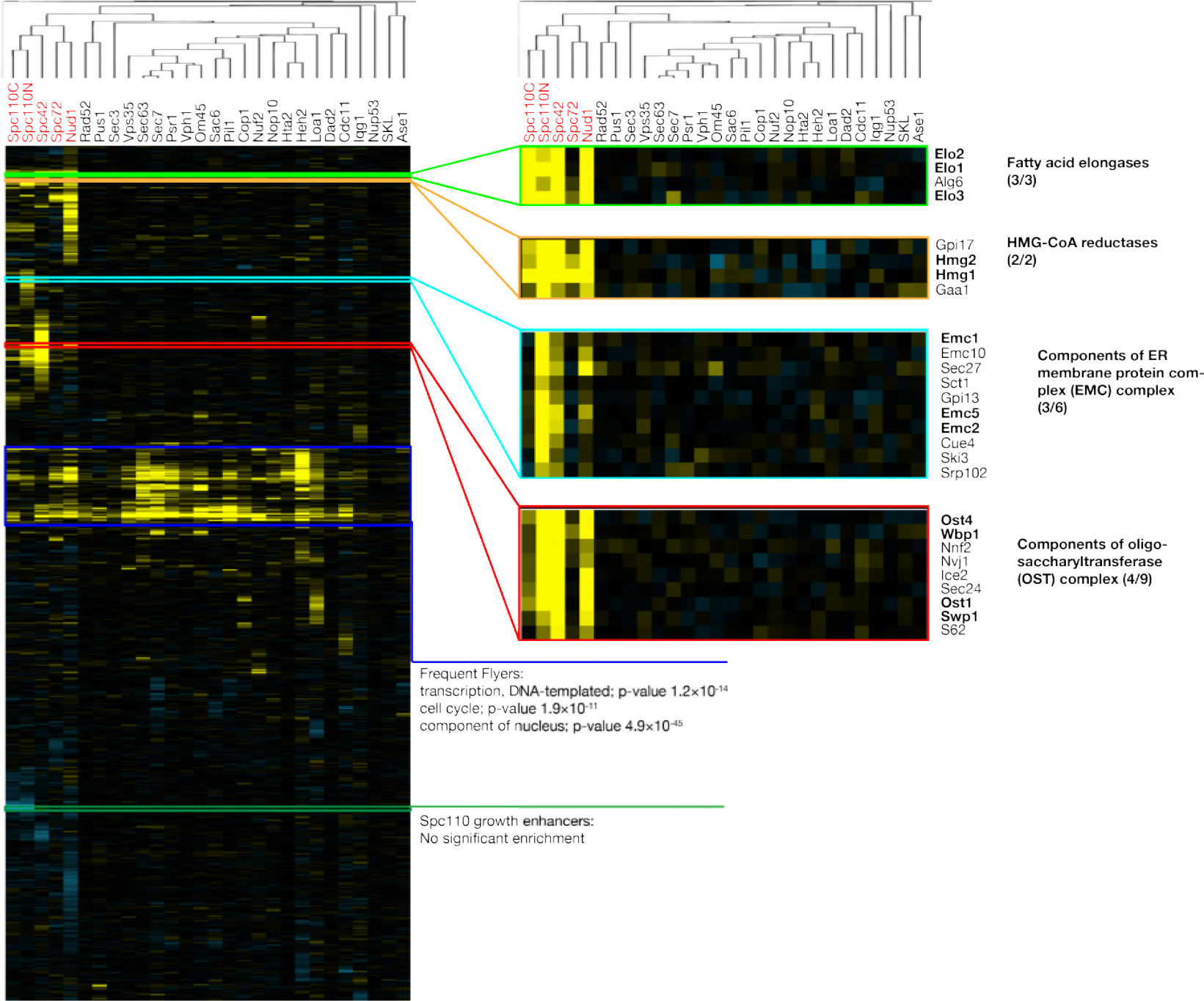
Cluster analysis of all 28 SPI screens used in this study. The data is clustered both vertically (by GFP strain) and horizontally (by screen, tree shown). The horizontal clustering tree shows that the SPB screen results are more similar to each other than the other screens. The vertical clustering identifies clusters of biologically related proteins with similar profiles of sensitivity to forcible relocalization.

In order to understand which kinds of proteins are sensitive to forced relocalization to each part of the SPB, we performed Gene Ontology (GO) analysis of the ranked LGRs for each of the screens using the GOrilla tool [Eden et al., 2009]. Heatmaps of significant enrichments are shown in Figure 5. The screens with Spc42, Spc110C and Spc110N fusions were all significantly enriched for proteins involved in lipid metabolic process and proteins from the ER. In particular, there was significant enrichment for proteins involved in biosynthesis of sterols, sphingolipids and those involved in fatty acid elongation (Supp. Figure 5). The position of the SPB, embedded within the nuclear membrane [Jaspersen and Winey, 2004], suggests that these growth defects may result from disregulation of nuclear membrane composition. Furthermore Witkin et al. [2010] found that deletion of *SPO7*, a regulator of phospholipid biosynthesis, could partially suppress the monopolar phenotype of mutations in *MPS3*, suggesting that the membrane environment can impact on SPB duplication. We also found that the screens with Nud1, Spc72 and Spc110N, the proteins located closest to the sites of microtubule nucleation, were enriched for proteins involved in the process of microtubule nucleation. These findings suggest that targeting these proteins artificially to the SPB can induce growth defects, possibly due to problems with spindle formation or nuclear positioning. An intriguing result is the finding that all screens, except Spc42, were enriched for proteins involved in chromosome segregation and components of the chromosome and kinetochore; there is evidence for example from yeast-two-hybrid screens that kinetochore proteins physically interact with SPB components [Wong et al., 2007]. It is worth noting that these phenotypes may simply represent disruption of these structures by removal of the protein to the SPB, although these proteins were not frequent flyers. Nud1 and Spc72 are thought to act as a signalling scaffold for proteins in the Mitotic Exit Network pathway [Scarfone and Piatti, 2015] and screens with these proteins were enriched for mitotic cell cycle proteins. Finally, we found that the Spc42 screen was enriched for proteins involved in nuclear pore organization as well as subunits of the nuclear pore. Intriguingly, some of these findings overlap with known genetic interactions, for example deletion of *NUP157* sup-presses the *spc42-11* mutation [Witkin et al., 2010], while we found that tethering Nup157 to Spc42 lead to a growth defect. Due to the proposed link between SPB duplication and insertion and the nuclear pore [Rüthnick and Schiebel, 2018] we investigated these results further.

**Figure 5:**
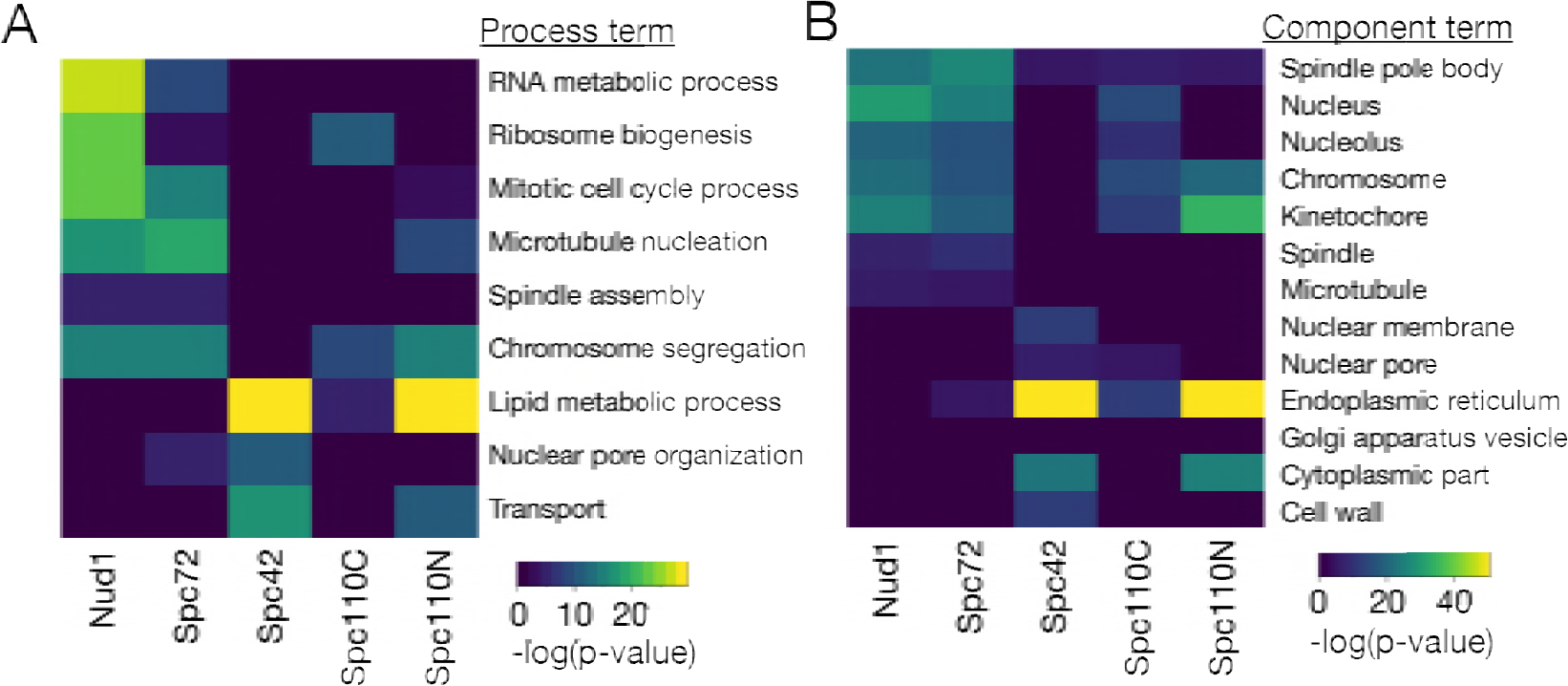
GO analysis of SPB SPI screens. A: Heatmap of process GO analysis, dark blue tiles represent no significant enrichment while the lighter colours represent significant enrichment, with warmer tones representing higher p-values. B: Heatmap of component GO analysis.

### 4.5 SPIs with the SPB lead to SPB overduplication

We investigated whether we could detect any SPB duplication phenotype caused by forcible localization of proteins to the SPB. We screened 80 query proteins that we suspected would cause defects in SPB duplication against the Spc42-GBP-RFP fusion. These proteins in-cluded proteins known to play a role in SPB duplication, such as the SPIN network and nuclear pore complex proteins as well as other transport proteins and hits from the screen that are as yet un-annotated. Cells were imaged using fluorescence microscopy and the number of SPBs, as approximated by the number of RFP foci, were counted. In particular, we searched for cells with 3 or more foci. In strains expressing membrane or pore proteins tagged with GFP we observed recruitment of Spc42-GBP-RFP to these regions, however small regions with relatively high RFP signal were observed and interpreted to represent SPBs. We found evidence of extra SPBs in eleven different strains (Table 1). In some cases, a single red focus was observed in large budded cells, however the slow-folding nature of RFP prevented us from ruling out the possibility of further SPBs that are unmarked by mature RFP. Screening cells directly from the SPI screen meant that limited number of cells were available to image in slow-growing SPI strains. Therefore, we directly transformed these strains, alongside the four members of the SPIN network, with the Spc42-GBP-RFP plasmid. We were able to establish colonies of all strains except Crm1-GFP. Using these strains we imaged larger quantities of these cells (Figure 6A). We detected extra red foci in each of these strains and quantified the proportion of cells expressing this phenotype (Figure 6B). Notably, we found that the strength of growth defect as measured by the LGR was not a strong indicator of the frequency of extra red foci, suggesting the growth defect does not arise entirely from this phenotype. Note that the protein denoted by its ORF, YJL021C, is included in these results however this ORF was determined to overlap *YJL020C* Brachat et al. [2003] meaning the GFP product in this strain is likely not a simple N-terminal fusion. Furthermore, the GFP strain shows a punctate fluorescent signal, meaning the extra red foci in these cells may represent relocalization of Spc42-GBP-RFP to YJL021C-GFP foci. These results suggest that the forced interaction of these proteins with the SPB results in aggregates of the Spc42 protein that may indicate extra SPBs.

**Figure 6:**
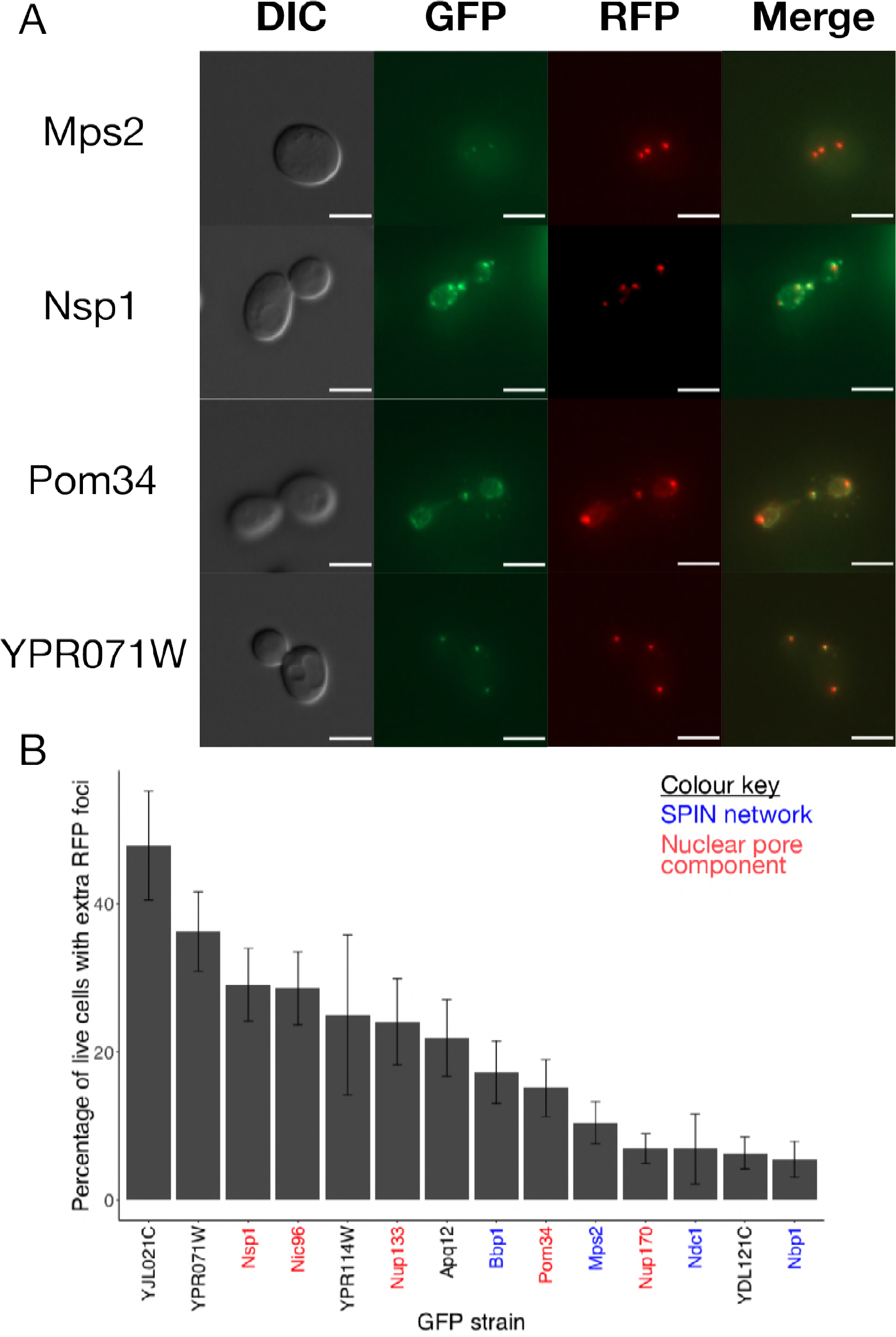
The plasmid encoding Spc42-GBP-RFP was directly transformed into the GFP strains identified in the screen for SPB number aberrations. A: Representative images of Mps2, Nsp1, Pom34 and YPR071W-GFP strains, each showing more than two RFP foci, interpreted as indicative of overduplication of SPBs. All scale bars are 5*μ*m. B:Quantification of the microscopy analysis, with key proteins highlighted, error bars show binomial standard error. Three images were captured for each strain and the percentage of living cells showing more than two RFP foci was calculated. Note that even relatively small percentages may be of interest as SPB overduplication is never observed in wild type cells.

**Table 1:** Proteins identified in the microscopy screen for proteins that induce extra SPBs when forcibly relocalised to the SPB. * YJL021C overlaps the originally identified YJL020C ORF and so has been merged into YJL020C [Brachat et al., 2003], however the GFP strain shows a punctate fluorescent signal.

| Protein | Database Location | Screen LGR | Retest LGR |
| --- | --- | --- | --- |
| Apq12 | ER | 1.58 | 1.46 |
| Crm1 | Nucleus | 1.23 | 0.54 |
| Nic96 | Nuclear Periphery | 1.06 | 1.20 |
| Nsp1 | Nuclear Periphery | 0.91 | 0.36 |
| Nup133 | Nuclear Periphery | 0.23 | 0.27 |
| Nup170 | Nuclear Periphery | 0.19 | 0.33 |
| Pom34 | Nuclear Periphery | 0.46 | 0.77 |
| YDL121C | ER | 2.40 | 1.83 |
| YJL021C* | Punctate | 0.74 | 0.95 |
| YPR071W | ER | 0.42 | 0.82 |
| YPR114W | ER | 2.67 | 2.01 |

**Table 2:** Table of plasmids

| Plasmid Name | Genotype | Selection |
| --- | --- | --- |
| pHT4 | *GBP* | Leu\Amp |
| pHT11 | *SPC42-GBP-RFP* | Leu\Amp |
| pHT297 | *SPC42* | Leu\Amp |
| pHT575 | *SPC110-GBP-RFP* | Leu\Amp |
| pHT576 | *GBP-RFP-SPC110* | Leu\Amp |
| pHT577 | *SPC110* | Leu\Amp |
| pHT584 | *NUD1-GBP-RFP* | Leu\Amp |
| pHT585 | *NUD1* | Leu\Amp |
| pHT615 | *SPC72* | Leu\Amp |
| pHT616 | *SPC72-GBP-RFP* | Leu\Amp |
| pHT706 | *HTB2-AZURITE* | Nat\Amp |

## 5 Discussion

### 5.1 Analysis of SPI screens

Z-transformations are a commonly used tool to analyse genome-wide screens, however their underlying assumption of normally distributed data means they can produce unreliable results, especially in screens identifying many hits. We developed empirical Bayes approach to address the shortcomings of Z-transformations. We developed tools including cutoffs based on the probability of inclusion which prove more effective in discriminating between screens with many hits. Notably, some hits in the Spc42 screen, such as Nup133 and Nup170, had LGRs which would have been considered insignificant according to the Z-score cutoff, but show a distinct phenotype. This shows that the lower cutoffs we propose can still be biologically relevant. Additionally, we developed a method to predict the validation rate of the screen with reasonable accuracy. These two metrics provide alternative viewpoints on the significance and strength of a given result within a screen. Furthermore, bimodal normal mixture models have 5 independent parameters, allowing for more effective parameterisation of the distribution than the 2 parameters of a single normal distribution. These parameters can be used to compare the distribution of results from different screens, providing a way to understand the differences between SPI screens. Previous analysis of cell-wide SPI screens concluded that only a small proportion of proteins are sensitive to forced relocalization however our re-analysis of this data and the inclusion of the SPB in this dataset suggests that some regions of the cell are far more sensitive to forced relocalization of proteins across the proteome. The empirical Bayes approach provides the most utility in the cases where Z-transformations are least appropriate: when analysing screens with large numbers of hits. However, we did find that when the fitted standard deviation of Component 2 was too large, the screens validated poorly and the mixture model was inaccurate.

Mixture models allow for a quick and easy way to effectively parameterise screening distribution data, however they cannot provide perfect prediction with imperfect data. If greater levels of precision or reproducibility are necessary, further modifications to the experimental procedures would be required. Zackrisson and colleagues found significant variation in growth rates of colonies across a single plate and recommend local normalisation of colony size to account for these effects to improve reproducibility [Zackrisson et al., 2016]. Baryshnikova and colleagues found that “batch” effects, caused by subtle differences in, for example, media composition or incubator temperature, between plates grown at different times, caused significant variation in colony sizes [Baryshnikova et al., 2010]. While all plates in a SPI screen are generally grown concurrently, the validation screens were performed afterwards, once analysis of the screen has been performed. This may explain the high FPRs in validation of some screens. Baryshnikova and colleagues propose using linear discriminant analysis to compute “batch signatures” which could be used to limit batch effects. Finally, the precision of measurements could be improved by a methodology that directly correlates growth measurement to rate, for example by calculating a growth curve using automated scanning of plates at regular intervals [Zackrisson et al., 2016].

### 5.2 SPB overduplication

Our current understanding of the SPB duplication cycle of *S. cerevisiae* is that alternating activities of the CDK-cyclin complex and Cdc14 phosphatase are responsible for once-per-cycle duplication of the SPB [Rüthnick and Schiebel, 2018]. This model suggests that SPB duplication is initiated while Cdc14 activity is at its peak but may not be completed until CDK activity increases and Cdc14 activity decreases later in the cell cycle. It has been proposed that the SPB satellite is inserted into the nuclear membrane using molecular machinery that is responsible for nuclear pore complex (NPC) insertion [Rüthnick and Schiebel, 2018]. We screened NPC proteins for SPIs with Spc42 and used fluorescence microscopy to estimate SPB number in these strains. We found evidence that the SPIN components, Bbp1 and Mps2, and to a lesser extent Nbp1 and Ndc1, induced formation of additional SPBs when forcibly recruited to the SPB. We also found that the NPC components Nsp1, Nic96, Nup133, Nup170 and Pom34 produced similar effects. It is interesting that the deletion of either of two of the genes coding for these proteins, *NIC96* and *POM34*, were identified as suppressors of SPB duplication defects caused by *mps3-1 spo7*Δ mutation [Witkin et al., 2010]. This finding suggests that in their wild type localization, these proteins inhibited SPB duplication, possibly by competing for binding partners, whereas our data suggests that when forced to the SPB, these proteins can induce the overduplication of SPBs. Additionally, we found evidence that the, as yet, unclassified proteins encoded by YJL021C, YPR071W, YPR114W and YDL121C as well as Apq12 similarly induce extra Spc42 foci indicative of SPB overduplication. There are several interpretations of these findings, which may apply to some or all of the phenotypes observed. Firstly, it is possible that the RFP foci observed may represent aggregates of Spc42-GBP-RFP that do not contain other SPB proteins or function as MTOCs. It is worth remarking that in a systematic study of localization of target and query proteins, a small proportion were found to localise to a region of the cell where neither would localise in wild type cells [Berry et al., 2016]. Secondly, it may be that forced recruitment of these proteins induce SPB overduplication through the documented SPB duplication pathway. This would require detachment of this process from the once-per-cycle regulation via CDK-cyclin and Cdc14. This could be explained if some aspects of this process were initiated by the presence of these proteins at the SPB, which were in turn induced by CDK-cyclin or Cdc14. Finally, it may be the case that targeting Spc42 to other structures in the cell, especially the NPC, can lead to the creation of de novo SPBs. The current model of SPB duplication suggests that SPBs assemble from a satellite formed of Spc42, Nud1, Cnm67 and Spc29 [Fu et al., 2015]. It may be that small amount of Spc42 are recruited to the NPC in these strains, and that these seed the creation of SPBs in a manner completely distinct from regular SPB duplication. Witkin et al. [2010] proposed an *MPS3* indepedent SPB duplication pathway and it may be this or some other pathway that is responsible for this phenotype.

Further work is required to distinguish these models, in particular, assessment of the foci for presence of other SPB proteins and functionality of the foci as MTOCs is required to confirm them as real SPBs. If SPBs are created de novo we would expect that these strains would lose the requirement for proteins with an essential role in SPB duplication, 452 such as Cdc31 [Rüthnick and Schiebel, 2016]. Many mutants have been identified that fail to duplicate their SPBs, for example the original MPS (Mono-Polar Spindle) genes [Winey et al., 1991] however there are fewer cases of genetic perturbations that lead to SPB overduplication. One example is the *sfi1-C4A* mutation [Avena et al., 2014], however this leads to SPB separation defects as well as SPB overduplication. The development of strains which reproducibly produce extra SPBs and multi-polar spindles will allow for the use of yeast models to explore the impact of these structures in cancers.

## 6 Acknowledgements

The authors acknowledge funding from the Francis Crick Institute, which receives its core funding from Cancer Research UK (FC001003), the UK Medical Research Council (FC001003), the Wellcome Trust (FC001003). We thank L Berry, G Ólafsson, P Bates, F Caudron, S Santos, W Taylor, U Eggert, J Diffley and the Genomics Equipment Park STP (The Francis Crick Institute). The authors declare no competing financial interests

## Supplementary Information - Figure 1

- fig1 - colocalization.xlsx
- fig1 - all screen LGRs.xlsx
- fig1 - validation.xlsx

## Supplementary Information - Figure 2

- fig2 - MM methods.pdf

## Supplementary Information - Figure 3

- fig3 - MMdata(GBlibrary).xlsx

## Supplementary Information - Figure 4

None.

## Supplementary Information - Figure 5

- fig5 - LipidGOanalysis.png
- fig5 - GOenrichment.xlsx

## Supplementary Information - Figure 6

- fig6 - allmicro1.png
- fig6 - allmicro2.png
- fig6 - microscopy screen.xslx
- fig6 - phenotype quantification.xlsx

## 8 Supplementary Figures

**Figure S5:**
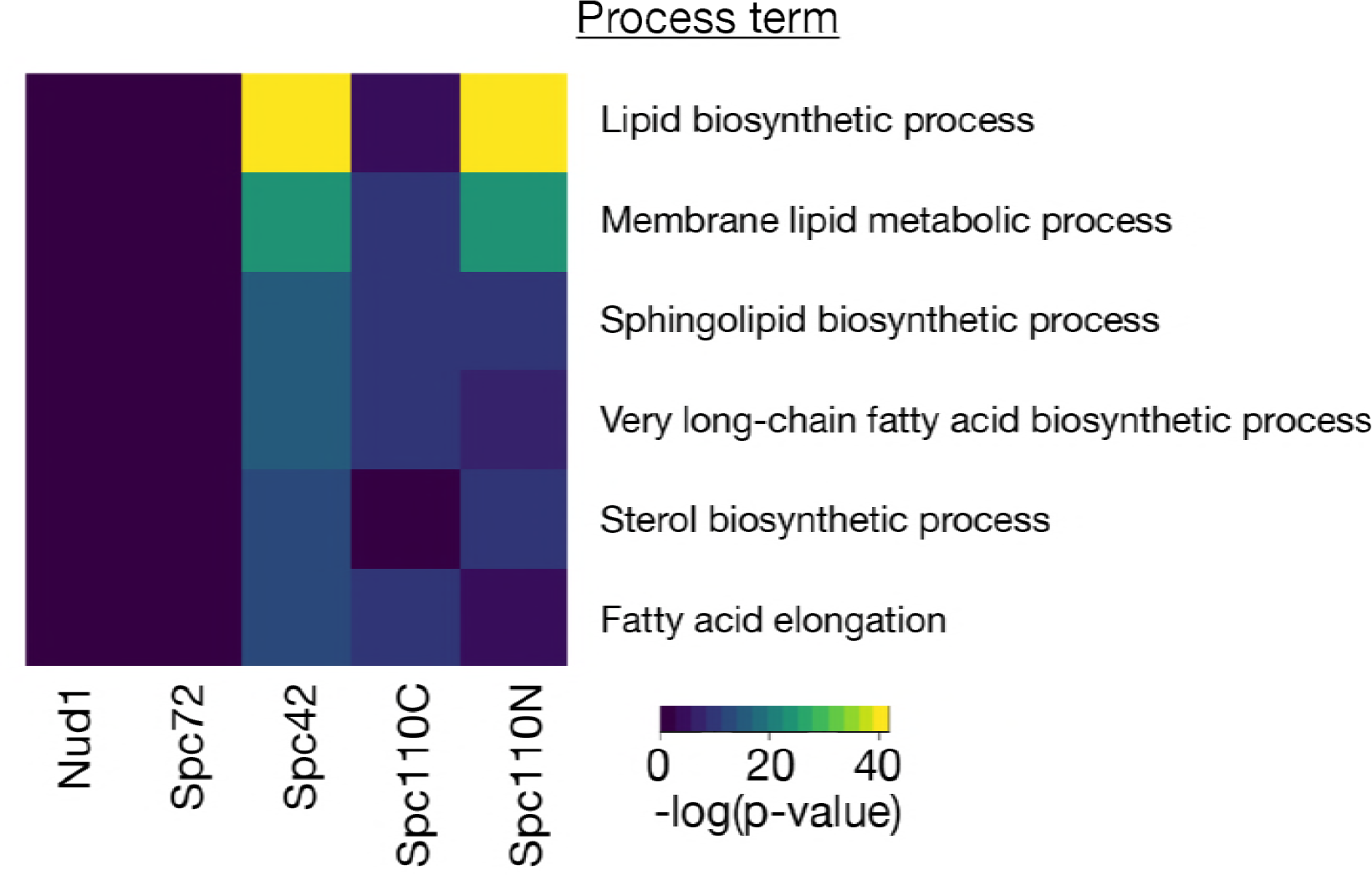
Heatmap of GO enrichment analysis for lipid process terms.

**Figure S6:**
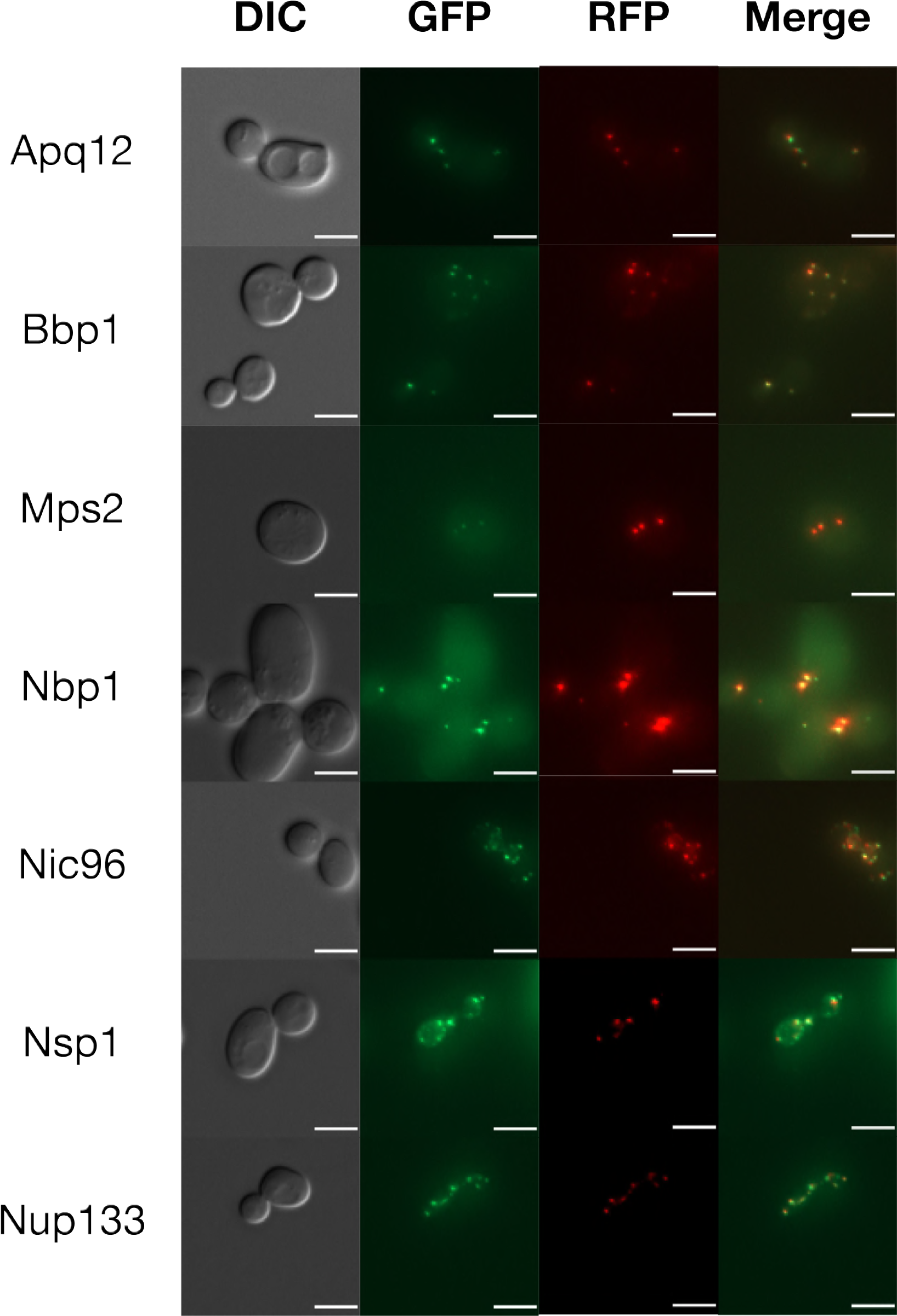
Representative images of each of the GFP strains investigated, each showing more than two RFP foci, interpreted as indicative of overduplication of SPBs. All scale bars are 5*μ*m.

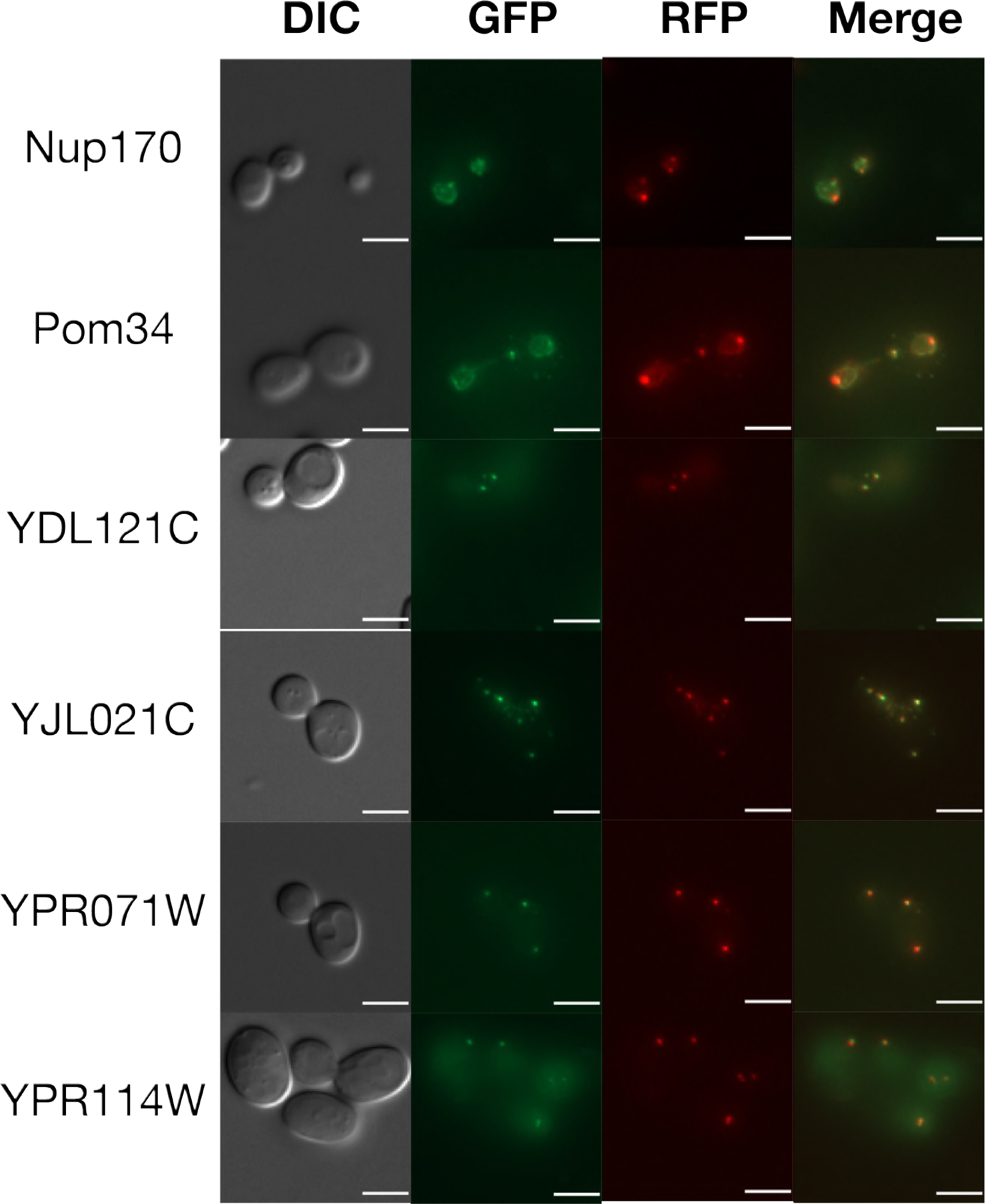

## Supplementary Methods

### Theory

Mixture models have been proposed as an alternative to calculating p-values based on the assumption that data is normally distributed Efron [2004] and have previously been used to analyse genome-wide datasets. The theory behind their use is that genome-wide screens are conducted in order to identify genes involved in a given process and that this divides the genome into two categories: those that are involved in this process (hits) and those that aren’t. Typically, non-hits will have a normal distribution centred around 0, due to variation caused by inherent noise in the system. In contrast, measurement of each of the hits can be thought of as a sample of a normal distribution with mean (and potentially variance) determined by the individual hit, the combination of these hits will form a distribution with properties that will depend on the biology of the screen. The aim of analysing genome-wide screen data is to distinguish these two categories. If there are few enough hits, they will simply form a tail at the edge of the distribution of non-hits and will not significantly effect the mean or standard deviation of the overall distribution. However, when there are significant numbers of hits, they will effect these summary statistics and a fitted normal distribution is unlikely to accurately reflect the real distribution of non-hits. This will render methods based on this approximation, such as the calculation of p-values and application of Z-transformations, inaccurate. The mixture model approach attempts to overcome this limitation by directly identifying the distribution of each of the two categories. Efron’s original method [Efron, 2004] involved fitting a normal component to the central peak of the data, representing non-hits, based on the shape of this peak. He then estimated the distribution of the hit peak from the difference between the overall distribution and the fitted null distribution. A limitation of this approach is that the null model is fitted to a relatively small region of the distribution of non-hits and furthermore, it gives no information about the distribution of the hits. In our approach, we fitted two normal modes to the data, using an Expectation-Maximization (ME) algorithm, which iteratively improves the fit of the model based on the likelihood of the generating the observed data from the given model. This means all of the data is used to fit the model and the end result is a parameterised model of the distribution of the hits which can be used to compare different genome-wide screens.

### Fitting

We fit two-peak normal mixture models to the smoothed LGR data for each of the screens, using the Mclust package Scrucca et al. [2016], which uses an ME algorithm to fit the model. The model fitting process yields 6 parameters: *ρ*_1_, *ρ*_2_, *μ*_1_, *μ*_2_, *σ*_1_, *σ*_2_ which fully define the mixture model. A table of all parameters of fitted models is included in the supplementary materials.

### Peak Identification

After fitting, we distinguished two types of fit: good fits that had two clearly defined distributions representing hits and non-hits; and poor fits where the distributions were not clearly defined. These poor fits were defined as those in which

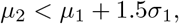

these screens were excluded from further analysis with mixture models. In the remaining 20 cases where the fit was good, we identified the “hit peak” as the peak shifted furthest to the right and the distribution of non-hits, or “central peak” as the leftmost distribution. We refer to these two components 40 of the distribution as *C*_1_ for the central peak and *C*_2_ for the hit peak. We can consider the genome-wide screen as a process for assigning LGRs to particular genes, the first step of this process is to decide whether the gene is a hit or not, which is a Bernouilli variable or weighted coin flip, where the probability of being a hit is given by *ρ*_2_. Then a gene *G*_*i*_ has identity *I*_*i*_ given by:

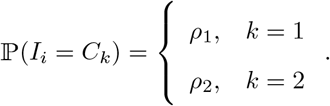

Once the identity is determined, the measured LGR, *LGR*_*i*_, is assigned as a normal variable distributed with mean and standard deviation *μ*_1_, *σ*_1_ or *μ*_2_, *σ*_2_ as determined by the category in which the gene was placed.

We wanted to define metrics to inform about the significance of results. In some cases we wish to draw a line that distinguishes LGRs from hits and non-hits and these metrics allow for such definitions. While cutoffs are a widely used tool and help to focus on significant results, they will always be to some extent arbitrary, as cases on the border may be placed either side by chance. On top of this, the strength of the interaction will vary depending on the particular genes, and depending on the application we may want only strong hits or we may want to include more subtle phenotypes. Therefore we propose different metrics to give a fuller picture of the data and so that a relevant metric can be chosen depending on context.

### p-value and Adjustments

The central peak of the distribution provides a natural null model for the data and this can be used to calculate a p-value for a given LGR, *x*:

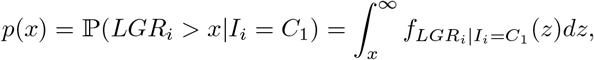

where *fX* (*x*) represents the probability distribution function of the random variable *X*. This value gives a measure of the probability that a given LGR would have been measured if the identity of gene *G*_*i*_ was the central peak *C*_1_. Genome-wide screens test multiple hypotheses so we may adjust the p-values to account for this, using for example either Bonferroni or FDR q-value adjustments [Benjamini and Hochberg, 1995]. A p-value of 0.05 is generally considered to be the cutoff for significance.

### Probability of Inclusion

As the intention of a genome-wide screen is to distinguish hits from non-hits, rather than considering the p-value we can consider the probability of inclusion in a given category. For a given LGR, *x*, the probability of inclusion in Component 2 is:

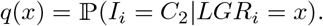

By Bayes’ theorem

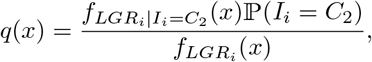

where 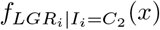 and 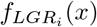 can be calculated from the fitted distributions. A sensible cutoff according to this approach is the point where a given gene is more likely to belong to Component 2 than Component 1, in other words *q*(*x*) = 0.5. We refer to this cutoff as *L*_*q*,0.5._

### Validation prediction

We validated our SPI screens against GFP-free controls, however this can be a time-consuming activity and so we developed analytical methods to predict the probability of validation. A strain is considered to be a validated hit if its retested LGR exceeds the mean plus two standard deviations of the LGRs of GFP-free controls on the plate. Note this is different to the methodology of Berry et al. [2016], in which the maximum LGR of the GFP-free controls was used as a cutoff. We define the probability of validation for a given LGR, *x* to be:

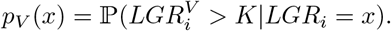

Using the law of total probability and conditioning on which of the categories gene *G*_*i*_ belongs to,

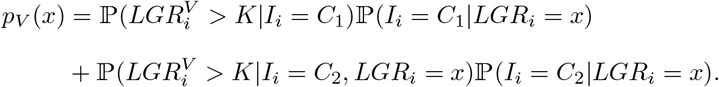

These values may all be simply calculated from the fitted mixture model, with the exception of 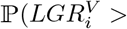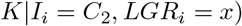. We assume that

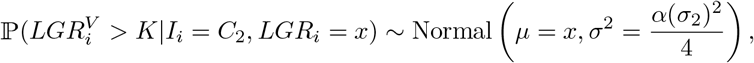

where *α* is a tunable parameter. We chose to centre the distribution on the original measurement of the LGR based on our observation that generally validation LGRs are similar to the genome-wide screen values. The variance of this distribution is not trivial to describe as it represents both noise in the system and batch effects. We chose to use 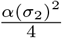, where the factor of four is derived from the higher density of colonies (16 rather than 4) used in the retest, and *α* is a tunable parameter representing batch effects. We found good accuracy using *α* = 4 and used this in all analysis.

We found that *pV* (*x*) performed well at predicting validation rate and FPR, with some exceptions (see main text). We propose that the curve *pV* (*x*) could be used as a tool when making decisions about how many results to validate in a genome-wide screen.

### Code accessibility

R scripts for data formatting and analysis are freely available at https://github.com/RowanHowell/data-analysis.

